# Genomic Diversity and Antimicrobial Resistance of *Vibrio cholerae* Isolates from Africa: A PulseNet Africa Initiative Using Nanopore Sequencing to Enhance Genomic Surveillance

**DOI:** 10.1101/2024.12.17.628868

**Authors:** Ebenezer Foster-Nyarko, Shola Able-Thomas, Nana Eghele Adade, Rexford Adade, Jean Claude Blessa Anne, Loretta Antwi, Yaya Bah, Gifty Boateng, Heather Carleton, David Chaima, Roma Chilengi, Kalpy Julien Coulibaly, Firehiwot Abera Derra, Dwayne Didon, Cheelo Dimuna, Mireille Dosso, Momodou M. Drammeh, Sana Ferjani, Kathryn E. Holt, Rohey Jatta, John Bosco Kalule, Abdoulie Kanteh, Hortense Faye Kette, Dam Khan, N’da Kouame Nazaire Kouadio, Christine Lee, Hamakwa Mantina, Gillan Mulenga, John Mwaba, Fatou Nyang, Godfred Owusu-Okyere, Jessica Rowland, Aissatou Seck, Abdul Karim Sesay, Anthony Smith, Peyton Smith, Djifahamaï Soma, Nomsa Tau, Pierrette Landrie Simo Tchuinte, Peggy-Estelle Maguiagueu Tientcheu, Chalwe Sokoni, Sabine N’dri Vakou, Delfino Vubil, PulseNet Africa

## Abstract

**Objectives:** *Vibrio cholerae* remains a significant public health threat in Africa, with antimicrobial resistance (AMR) complicating treatment. This study leverages whole-genome sequencing (WGS) of *V. cholerae* isolates from Côte d’Ivoire, Ghana, Zambia and South Africa to assess genomic diversity, AMR profiles, and virulence, demonstrating the utility of WGS for enhanced surveillance within the PulseNet Africa network.

**Methods:** We analysed *Vibrio* isolates from clinical and environmental sources (2010–2024) using Oxford Nanopore sequencing and hybracter assembly. Phylogenetic analysis, multilocus sequence typing (MLST), virulence and AMR gene detection were performed using Terra, Pathogenwatch, and Cloud Infrastructure for Microbial Bioinformatics (CLMB) platforms, with comparisons against 88 global reference genomes for broader genomic context.

**Results:** Of 79 high-quality assemblies, 67 were confirmed as *V. cholerae*, with serogroup O1 accounting for the majority (43/67, 67%). ST69 accounted for 60% (40/67) of isolates, with eight sequence types identified overall. Thirty-seven isolates formed novel sub-clades within AFR12 and AFR15 O1 lineages, suggesting local clonal expansions. AMR gene analysis revealed high resistance to trimethoprim (96%) and quinolones (83%), while resistance to azithromycin, rifampicin, and tetracycline remained low (≤7%). A significant proportion of the serogroup O1 isolates (41/43, 95%) harboured resistance genes in at least three antibiotic classes.

**Conclusions:** This study highlights significant genetic diversity and AMR prevalence in African *V. cholerae* isolates, with expanding AFR12 and AFR15 clades in the region. The widespread resistance to trimethoprim and quinolones raises concerns for treatment efficacy, although azithromycin and tetracycline remain viable options. WGS enables precise identification of species and genotyping, reinforcing PulseNet Africa’s pivotal role in advancing genomic surveillance and enabling timely public health responses to cholera outbreaks.

**Data summary:** All supporting data and protocols have been provided within the article or as supplementary data files. The ONT reads have been deposited under BioProject accession PRJNA1192988, while the high-quality *Vibrio* spp. assemblies have been shared via figshare (Foster-Nyarko, Ebenezer (2024). Genomic Diversity and Antimicrobial Resistance of Vibrio spp. Isolates from Africa: A PulseNet Africa Initiative Using Nanopore Sequencing to Enhance Genomic Surveillance. figshare. Dataset. https://doi.org/10.6084/m9.figshare.27941376.v1). Individual accession numbers for these reads and Biosample IDs are provided in **File S2,** available with the online version of this article. The accession numbers for the 88 reference genome assemblies included in our analysis are also provided in **File S3**.

**Impact statement:** Cholera remains a significant public health challenge in Africa, disproportionately affecting the region due to the ongoing transmission of *Vibrio cholerae* O1 and the emergence of antimicrobial resistance (AMR). This study demonstrates the utility of Oxford Nanopore Technology (ONT) sequencing in providing high-resolution insights into the genomic diversity, transmission dynamics, and AMR profiles of *V. cholerae* isolates across Africa. By generating and analysing whole-genome sequences, we identified novel sublineages, high prevalence rates of AMR genes, and virulence traits critical to cholera pathogenesis. These findings contribute to a deeper understanding of the epidemiology and evolution of *V. cholerae* in Africa, informing targeted intervention strategies.

Furthermore, the study highlights the growing threat posed by AMR among *V. cholerae* isolates, including resistance to key therapeutic antibiotics, such as quinolones and trimethoprim, which could undermine current treatment protocols. Despite this, the absence of resistance to azithromycin and rifampicin among the O1 isolates suggests these drugs may remain viable treatment options, offering a critical avenue for preserving treatment efficacy.

This research also underscores the importance of sustained genomic surveillance, capacity building, and regional collaboration to mitigate the public health impact of cholera and other foodborne pathogens. By leveraging WGS technologies and training initiatives, such as the PulseNet Africa genomics workshop, this study provides a framework for strengthening regional capacities to detect, monitor, and respond to cholera outbreaks and the spread of AMR. These efforts align with the African Union and Africa CDC’s strategic priorities on health security and AMR, contributing to improved public health systems and cholera control across the continent.

## Introduction

Cholera, caused by toxigenic strains of *Vibrio cholerae* O1 and O139, presents with mild to potentially fatal acute watery diarrhoea (1–3). The current seventh cholera pandemic, primarily due to the O1 biotype El Tor, disproportionately affects Africa, which bears the majority of the global disease burden (2). Despite its longstanding presence since resurging in 1970, the transmission mechanisms of *V. cholerae* in Africa remain poorly understood. This knowledge gap is exacerbated by shifts in antimicrobial resistance (AMR) patterns, moving from traditional antibiotics like streptomycin, chloramphenicol, furazolidone, and trimethoprim-sulfamethoxazole to tetracyclines, fluoroquinolones, and macrolides, climate change, and increased mobility of people, globally (3–7).

Recent genomic analyses have revealed multiple introductions of the *V. cholerae* serogroup O1 biotype El Tor strain from Asia to Africa, with at least 13 distinct sublineages identified since 1970, indicating a continued pattern of introduction from Asia (8–11). Understanding the genomic epidemiology and AMR profiles of *V. cholerae* in Africa is essential for developing effective surveillance and intervention strategies. Conventional methods like culture and antimicrobial susceptibility testing, PCR, and point-of-care testing have been instrumental in providing real-time information on circulating serotypes and AMR patterns in endemic countries in sub-Saharan Africa. However, these methods offer limited resolution and are insufficient for tracking the spread of resistance and understanding the evolutionary dynamics of circulating *V. cholerae* clones. Whole-genome sequencing (WGS) technologies, including Illumina and Oxford Nanopore Technology (ONT), offer improved resolution for characterising outbreak isolates and tracing transmission pathways. By providing comprehensive genomic data, WGS enables a deeper understanding of *V. cholerae* persistence, transmission, and AMR evolution in endemic regions. Such insights are vital for informing more effective cholera control strategies and public health responses.

Given its global distribution, international transmission, and endemicity across most of sub-Saharan Africa, *V. cholerae* has been a priority pathogen for PulseNet Africa since the network’s inception in 2010. PulseNet Africa is a regional collaboration of public health laboratories dedicated to tracking food-and water-borne disease outbreaks, including AMR. Alongside *V. cholerae*, PulseNet Africa monitors *Salmonella enterica*, diarrhoeagenic *Escherichia coli*, *Shigella* spp., *Campylobacter* spp., and *Listeria monocytogenes* across 19 African countries (**Figure 1**). It operates within the global PulseNet International network, collaboratively connecting over 108 countries in eight regions to monitor and control foodborne disease outbreaks.

**Figure 1.**
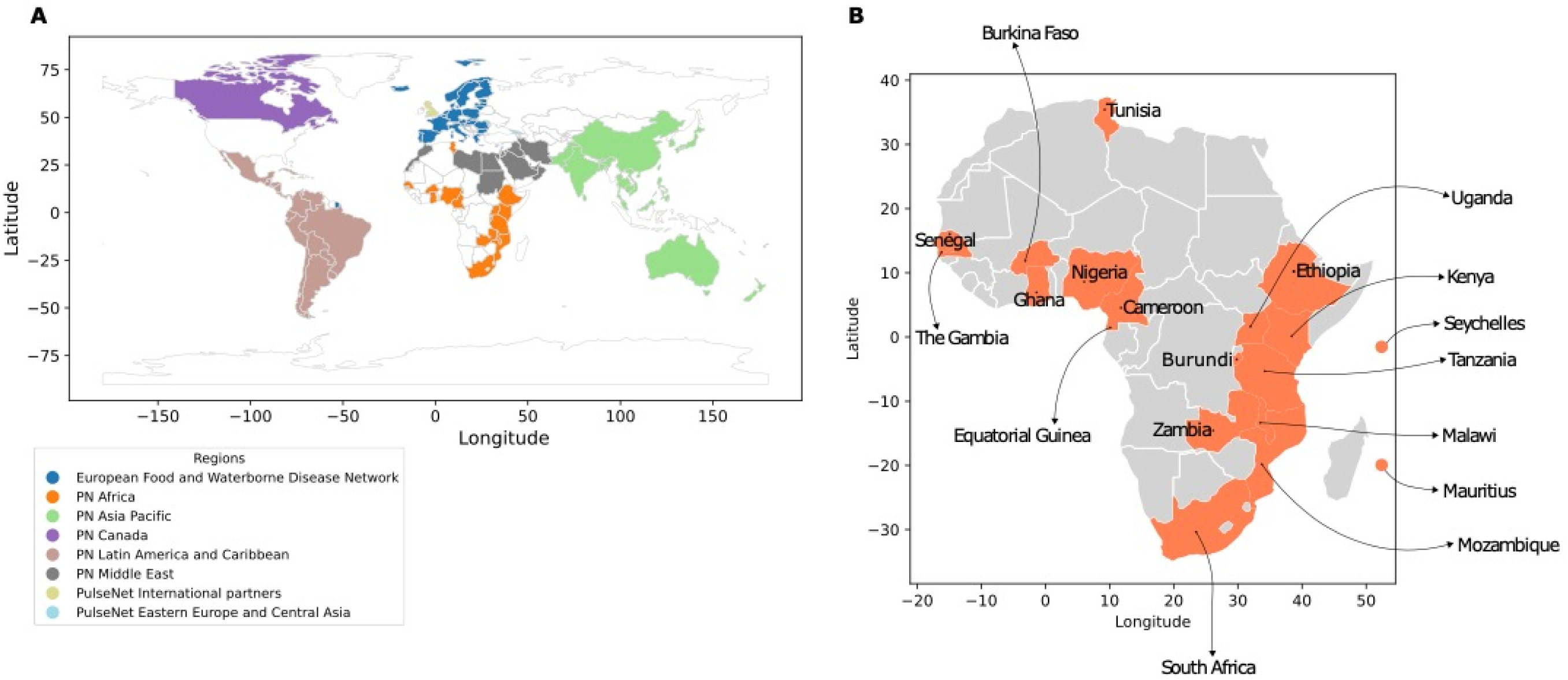
PulseNet International Regions and PulseNet Africa Member Countries. (**A**) A depiction of PulseNet International Regions, colour coded as follows: blue, European Food and Waterborne Disease Network; orange: PulseNet Africa; green: PulseNet Asia Pacific; purple: PulseNet Canada; brown: PulseNet Latin America and Caribbean; grey: PN Middle East; light yellow: PulseNet International Partners; light cyan: PulseNet Eastern Europe and Central Asia. The participating countries representing Africa, Asia Pacific, Eastern Europe and Central Asia, the European Food and Waterborne Disease Network, Latin America and Caribbean, Middle East and PulseNet International Partners are denoted in the corresponding colour-coded bubbles. (**B**) This panel zooms in on the PN Africa region and highlights its member countries.

The World Health Organization (WHO) recognises the critical role of WGS in enhancing global public health strategies, particularly for foodborne pathogens and AMR surveillance (12–14). In the context of foodborne pathogen surveillance, the WHO highlights the utility of WGS in outbreak investigations, response efforts, and routine monitoring. WGS facilitates precise pathogen identification, source tracing, and an improved understanding of genetic diversity. This precision is especially significant within a One Health framework, where human, animal, and environmental health are interconnected. By enabling more effective tracking of foodborne diseases, WGS strengthens public health interventions and supports global health security (12, 13).

Additionally, WGS offers invaluable insights into resistance mechanisms, as well as the emergence and spread of AMR, informing evidence-based policy development for AMR control (13). As countries advance toward cholera elimination, WGS analyses can be particularly impactful, distinguishing between rare, newly detected cases and those arising from previously circulating endemic strains. Thus, the integration of WGS into national and international surveillance systems is essential for enhancing precision, enabling swift outbreak responses, and promoting accurate cross-border data sharing.

Recently, PulseNet International adopted whole-genome sequencing in place of Pulsed-Field Gel Electrophoresis (PFGE) and multiple-locus variable number tandem repeat analysis (MLVA) for molecular sub-typing, owing to the superior resolution of WGS, which has substantially enhanced genomic epidemiological surveillance and tracking of AMR genes across bacterial populations (15–18). As molecular capacity continues to expand across the African continent, the implementation of standardised genomic surveillance for high-priority pathogens has become increasingly feasible (19). Additionally, the upcoming discontinuation of support for BioNumerics—the primary software for analysing data generated through PFGE, MLVA, and WGS—beyond 2024 presents a timely opportunity to evaluate and adopt alternative analysis tools that can address the diverse needs of the network.

To strengthen the genomic sequencing capabilities of participating laboratories in PulseNet Africa, we organised a hands-on genomics workshop focusing on ONT sequencing in collaboration with the Medical Research Council Unit The Gambia at London School of Hygiene and Tropical Medicine (MRCG at LSHTM), Theiagen Genomics LLC, the Association for Public Health Laboratories (USA), and the US Centers for Disease Control and Prevention (CDC). This workshop, held in July 2024 at the MRCG at LSHTM, provided both wet lab training on ONT sequencing and dry lab training for genomic data analysis using the Terra platform. Participants brought DNA from archived, unsequenced isolates identified as *V. cholerae* from outbreaks or routine surveillance in their countries. These samples were used to train participants on PulseNet International’s protocols for ONT sequencing of food-and water-borne pathogens. The workshop culminated in collaborative data analysis, where participants interpreted the genomic data to gain insights into the transmission dynamics and AMR profiles of *V. cholerae* across Africa. We hypothesised that *V. cholerae* isolates from various African regions would exhibit distinct transmission events, lineages, and AMR profiles.

Our efforts align with the African Union and Africa CDC’s strategic priorities on health security and AMR. By elucidating the genomic diversity, evolution, and AMR dissemination of *V. cholerae* in Africa, this study provides crucial insights into the regional and global emergence of multi-drug-resistant strains. The genomics workshop and this study represent significant steps toward enhancing the capacity for genomic sequencing of *V. cholerae* and other critical pathogens within Africa. Additionally, establishing a skilled professional network will help foster ongoing collaboration and support, potentially contributing to long-term improvements in public health systems across the continent.

## Methods

### Archived V. cholerae isolates analysed in this study

We generated WGS data from 104 archived isolates initially identified as *V. cholerae* by conventional microbiological methods (**File S1**) from four countries: Côte d’Ivoire (n=50), Ghana (n=19), Zambia (n=29) and South Africa (n=6), between 2010 and 2024 (**Figure 2A**). These isolates represented unsequenced archived isolates derived from outbreaks or environmental surveillance (**File S1**). Metadata associated with the isolates included the year of collection and geographical information (i.e., patient residential province or city) (**File S2**).

**Figure 2.**
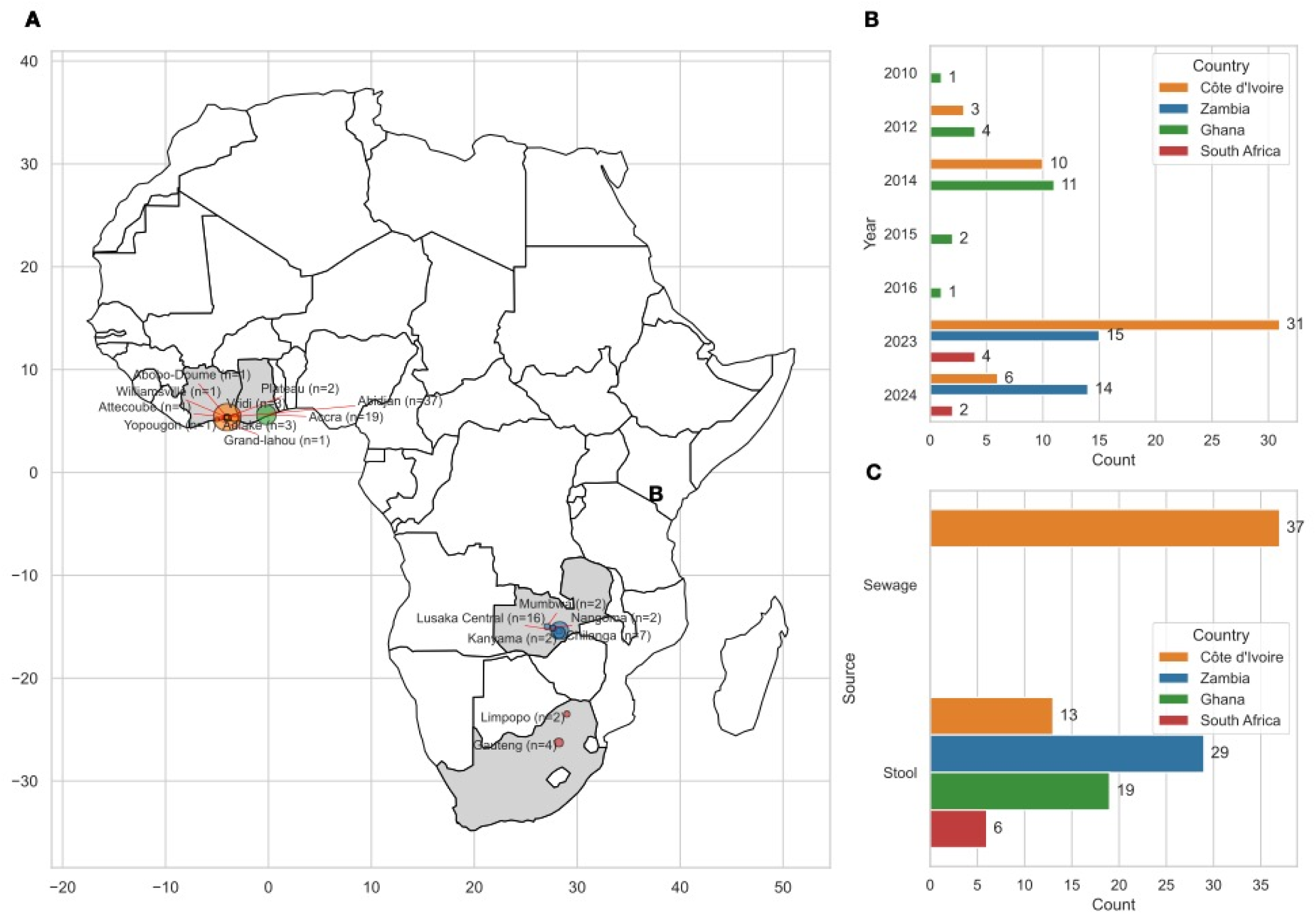
Geographical distribution, collection year, and sample source of the isolates analysed in this study. (**A**) Geographical distribution of the isolates analysed in this study. The map highlights the locations where the samples were collected. (**B**) The number of isolates collected per year is displayed, showing the temporal distribution of sample collection across different years per sampling site, as depicted in the legend. (**C**) The sample sources are categorised and plotted to represent the proportion of samples collected from outbreak (Stool) or non-outbreak (Sewage) situations.

### Global context isolates

To determine where our isolates sit within the global phylogeny, we included publicly available sequence data from 88 global reference strains of *V. cholerae* (**File S3**), representing the known diversity of *V. cholerae* serogroup O1 (8–11, 20–22), collected between 1970 and 2023.

### DNA extractions and ONT sequencing

DNA extractions were performed in PulseNet participating laboratories using standard protocols specific to each laboratory. In Zambia, extractions utilised the Invitek DNA extraction kit (Invitek Diagnostics, Germany), while isolates from Ghana were extracted using the Qiagen DNeasy blood and tissue kit (Qiagen Johannesburg). Nucleic acid extractions in Côte d’Ivoire were performed with the Spin-X viral DNA/RNA extraction kit (SD Biosensor, Republic of Korea), and in South Africa, the QIAmp DNA mini kit (Qiagen Johannesburg) was used. Library preparation followed PulseNet International’s standard operating procedures using ONT’s rapid barcoding kit.

All isolates, except those from South Africa, were sequenced during the training workshop at MRCG at LSHTM. The libraries were loaded onto 11 MinION R10 flowcells and sequenced on the GridION, Mk1cs, or Mk1bs platforms. In South Africa, sequencing was conducted at the National Institute for Communicable Diseases (NICD), and the ONT reads were basecalled using the same protocol employed at MRCG at LSHTM. The sequences from all locations were subsequently combined for analysis.

The basecalling utilised ONT’s basecaller, Dorado v0.7.2, with the SUPer-accuracy model (SUP) v4.3.0, via the high-performance computing clusters at MRCG at LSHTM or NICD. Subsequent bioinformatic analyses were conducted on the Terra platform (https://terra.bio/), the Cloud Infrastructure for Microbial Bioinformatics (CLIMB) (23) and Pathogenwatch (24).

### Bioinformatics analysis

The analysis on Terra utilised the TheiaProk_ONT_PHB v.2.1.0 and TheiaProk_FASTA_PHB v2.2.0 workflows. Initially, reads were quality-checked with FastQC (25). Assemblies were then generated on CLIMB using hybracter v0.7.3 (26) via CLIMB (23). The hybracter assemblies were then uploaded to Terra, where TheiaProk_FASTA_PHB v2.2.0 was deployed to characterise the genome assemblies. This included speciation with Gambit v1.0.0 (database v2.0.0- 20240628.gdb) (27), identification of *<u>omp</u>*W and t*ox*R loci and *Vibrio* characterisation using Abricate v1.0.1 (28), multilocus sequence typing using the mlst software v2.23.0 (https://github.com/tseemann/mlst), and plasmid identification with PlasmidFinder v2.1.6 (29).

Next, we uploaded the assemblies to Pathogenwatch (www.pathogen.watch), where a collection was created and neighbour-joining trees reconstructed using a concatenated alignment of 2,7336 genes which constitute the core gene library for *V. cholerae* in Pathogenwatch. PopPUNK (30) was deployed to determine the placement of our study isolates as belonging to the current pandemic lineage using the PopPUNK clusters implemented within Vibriowatch (24). PopPUNK classifies user-uploaded sequences to one of 41 stable clusters corresponding to the previously named *V. cholerae* lineages. For example, cluster 1 denotes isolates belonging to the current pandemic lineage (7PET), clusters 2 and 7 are part of Lineage 3b, while clusters 75 and 76 encompass strains that are part of ELA-1 (24). If a novel lineage is encountered, PopPUNK assigns cluster VC1870. For ONT assemblies, it is imperative to ascertain that the isolate truly belongs to a novel lineage and that sequencing errors are not hampering the PopPUNK analysis. Consequently, for each study isolate typed as VC1780 by PopPUNK or not belonging to previously characterised lineages, we created a Vibriowatch collection containing the study isolate and some contextual isolates from the Chun et al. (2009) collection (31), available in Pathogenwatch (24). This collection encompasses isolates from the 7PET lineage and the other known *V. cholerae* lineages (31). The neighbour-joining trees, generated by Vibriowatch, were then inspected to determine the clades the isolate fell into. Additionally, Pathogenwatch calls AMR genes using an in-house AMR search tool by scanning against a curated AMR gene library hosted on Pathogenwatch.

To contextualise the AMR genes and mutations found among the *V. cholerae* isolates against what pertains to the global *V. cholerae* O1 population, we searched for all *V. cholerae* O1 isolates in Pathogenwatch and downloaded the associated AMR genotype predictions and analysed the resulting data frame using Python’s pandas library v2.0.3. via Jupyter Notebook.

We then combined the *V. cholerae* ST69 sequences generated in this study with 88 publicly available representative sequences from the known 7PET *V. cholerae* sequences and performed a whole genome alignment of assemblies using the alignment-free mode implemented in the Split K-mer Analysis (SKA2) v0.3.7 (32)—a rapid and accurate approach which utilises k-mers to detect variation between samples. Single nucleotide polymorphisms (SNPs) detected by SKA have been shown to be comparable to those determined by read alignment-based methods such as Snippy (32–34). A maximum likelihood phylogeny was inferred from the resulting SNP alignment using RAxML v8.2.12 (35) with the generalised time reversible (GTR) substitution model and 1000 bootstrap replicates. The phylogenetic tree was visualised using FigTree v1.4.4 and annotated in R v4.1.0 (2021-05-18) with the ggtree package v3.0.4 and Adobe Illustrator v28.0.

## Results

In total, we sequenced 67 clinical and 37 environmental isolates previously identified as *V. cholerae*. The environmental isolates were collected exclusively from sewage samples in Abidjan, Côte d’Ivoire, between 2023 and 2024. For the clinical isolates, samples from Côte d’Ivoire were spread over seven towns or cities; those from Zambia originated from five sub-counties, while those from Ghana were sourced from the nation’s capital, Accra. The isolates from South Africa were derived from Limpopo and Gauteng (**Figure 2B-C and File S2**).

Since ONT sequencing for *Vibrio* is new to PulseNet International, and QC thresholds for ONT data are not yet established, we aimed to assess the assembler’s performance at varying read depths. We initially set a minimum read threshold of 5,000 reads, independent of sequencing depth, hypothesising that fewer reads would be sufficient for high coverage due to the longer ONT read lengths. All samples met this threshold and were assembled using Hybracter (26). The resulting assemblies were then examined with Bandage, which showed that samples with read coverage below 20x generally produced highly fragmented assemblies (>10 contigs) and low assembly coverage (<20x mean depth), except for three samples (CO42-15, CIV34, and ZA29) with read coverages of 8x, 13x, and 18x, respectively, which yielded fully circularised chromosomes. Therefore, in the downstream analysis, we only included samples with fully assembled chromosomes or a mean read depth and assembly coverage of ≥20x, excluding those below this threshold. A total of 25 samples, comprising 19 clinical and six environmental samples, fell below this threshold and were excluded, leaving a final dataset of 79 isolates. Detailed read and assembly metrics are presented in **Figure S1 and File S2**.

Of the 25 samples excluded from the final dataset due to insufficient read depth or assembly quality, 19 were stool samples, and 6 were from sewage. Despite not meeting the QC thresholds, some samples could still be speciated: 17 were identified as *V. cholerae*, 1 as *Aeromonas caviae*, and 1 as *Pseudomonas aeruginosa*, while 3 samples remained unidentified, and 3 failed to assemble altogether. Geographically, these excluded samples were distributed between Zambia (11), Côte d’Ivoire (11) and South Africa (3).

Of the 79 high-quality assemblies that constitute the final dataset, speciation with Gambit (27) and Pathogenwatch’s Speciator (36) were in agreement and revealed 67/79 (85%) isolates as *V. cholerae*, 2/79 (3%) each as *V. fluvialis* and *V. navarrensis*, and 1/79 (1%) as *V. furnissii.* The *V. furnissii* isolate was recovered from a clinical (stool) specimen in Vridi, Côte d’Ivoire, while the *V. fluvialis* and *V. navarrensis* isolates were all sourced from sewage samples in Abidjan (**File S2**). All isolates confirmed by WGS as non-cholerae *Vibrio* spp. resulted in complete circularised assemblies.

The remainder of the isolates typed as non-*Vibrio* species, as follows: *Enterobacter hormaechei* (4/79, 5%); *Aeromonas enteropelogenes* (2/79, 3%), and *Klebsiella quasipneumoniae* (1/79, 1%).

The 67 isolates confirmed by WGS as *V. cholerae* comprised n=46 from clinical (outbreak) cases (stool) and n=21 sewage isolates. Of these, 46 had two completely circularised chromosomes each, comprising n=30 clinical isolates and n=16 from sewage. The assembly metrics are are depicted in **Figure S1.**

The individual sample accessions for the high-quality *V. cholerae* isolates are provided in **File S2**, and the *Vibrio* spp. assemblies shared via figshare (https://doi.org/10.6084/m9.figshare.27941376.v1). The rest of the results focus on the *V. cholerae* isolates.

### Phylogenetic diversity of the study strains

We recovered eight sequence types overall, with ST69 accounting for the majority (40/67, 60%), followed by non-typeable (12/67, 18%), ST85 and ST75-2LV (2/67, 3% each), ST289, ST832, ST357, ST366 and ST69-4LV (1/67, 2% each). Six isolates were assigned one or more novel alleles out of the seven housekeeping genes utilised in the MLST typing, indicating potentially novel STs.

Most (43/46, 93%) of the clinical isolate genomes were identified as *V. cholerae* serogroup O1 and biotype El Tor, consistent with the serological results. The majority of *V. cholerae* O1 isolates belonged to ST69 or its close variant ST69-4LV and were assigned to the VC1 cluster by PopPUNK, indicating their association with the current pandemic lineage. Notably, two exceptions were observed: the ST75-2LV isolates from South Africa, which clustered within the pre-7PET clade (**Figure S2**). This clade is thought to represent close relatives of the most recent common ancestor of the 7PET clade (24). The nearest relative to the two ST75-2LV strains was isolate 2740-80, a US Gulf Coast *V. cholerae* O1 clone previously associated with the 2018– 2020 cholera outbreak in South Africa (37). In addition, two non-O1 clinical isolates (‘CIV50’ and ‘CIV55’), collected from Abobo-Doumé and Williamsville, Côte d’Ivoire, respectively, were identified as belonging to ST85 and were assigned to the VC995 cluster. A single environmental isolate from ST366 was assigned to VC506. The remaining non-O1 environmental isolates were grouped into VC1870 by PopPUNK, further underscoring their genomic diversity.

The phylogenetic analysis with representative global reference strains revealed distinct sub-lineages arising from local clonal expansions of AFR9, AFR12 and AFR15 among the *V. cholerae* O1 ST69 strains. Specifically, isolates ‘17-14’ and ‘C413-14’, collected in Accra in 2014, along with isolate CIV57 derived from Adiaké, Côte d’Ivoire, in 2012, clustered with reference genomes belonging to T9/AFR9, indicating genetic similarity to these known *V. cholerae* O1 strains. Likewise, n=22 isolates from Ghana (2010–2015) and Côte d’Ivoire (2012 – 2014) formed novel sub-clades within the AFR12 lineage (**Figure 3**), indicative of local clonal expansions of this lineage in these two countries. This sub-lineage was initially identified in Togo and thought to have been directly imported from South Asia (11). Similarly, the O1 isolates collected from central Lusaka, Nangoma and Kanyama in Zambia between 2023 and 2024 formed novel sub-clusters within the recently defined sub-lineage AFR15, which caused a large outbreak in South Africa between 2022 and 2023 (8), suggesting potential uncharacterised clonal expansions of this lineage within the *V. cholerae* O1 serogroup and underscoring the genetic diversity among our study isolates.

**Figure 3.**
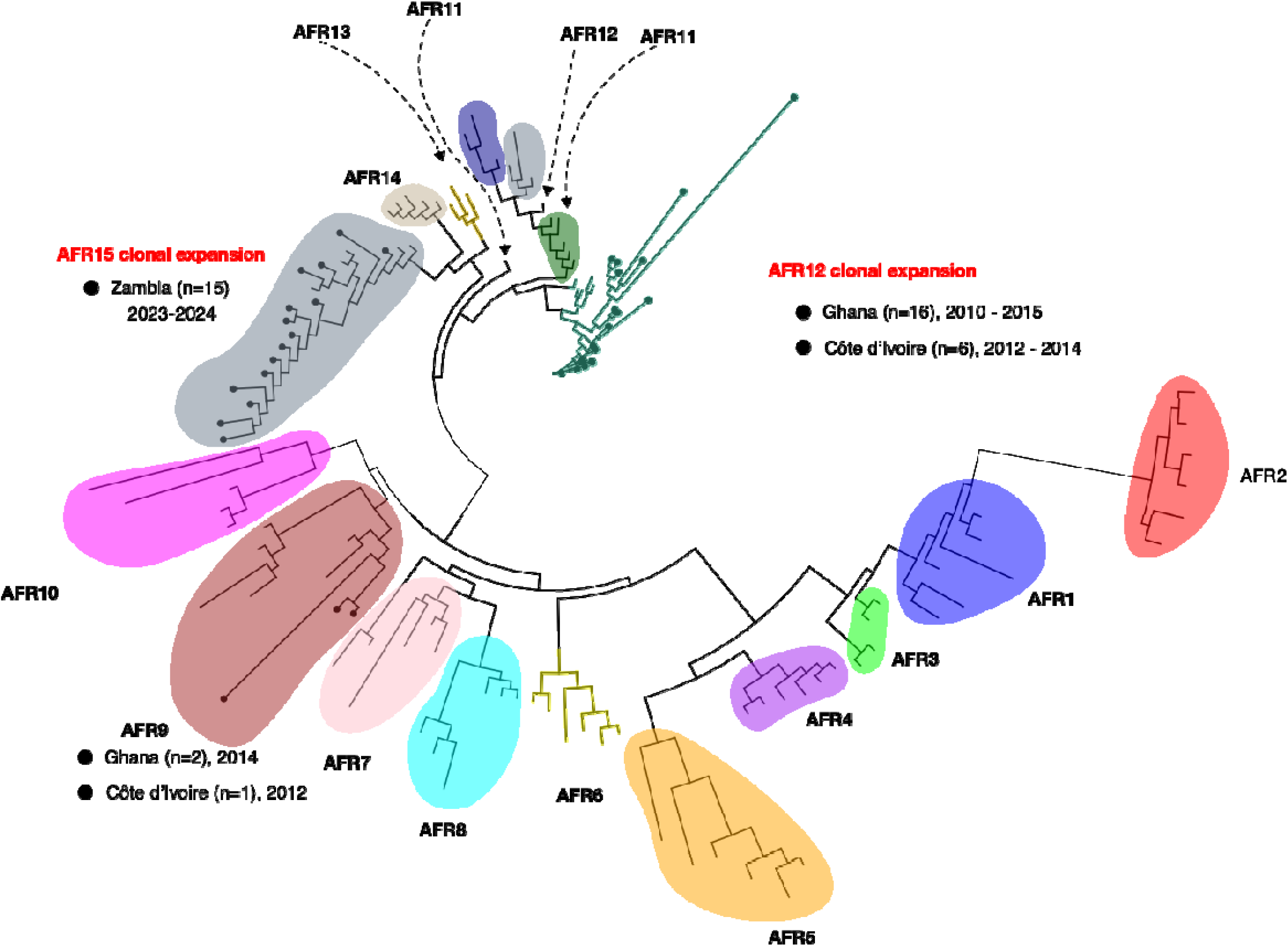
A Maximum-Likelihood tree depicting the population structure of the *V. cholerae* ST69 O1 study isolates (n=40), along with 88 publicly available genomes representing the known diversity of *V. cholerae* O1. The phylogenetic tree was reconstructed using RAxML v8.2.12 (Reference #35) with 1000 bootstrap replicates, based on a reference-free whole genome alignment generated with SKA2 (Reference #32). The branch lengths indicate genetic distances between samples. The clades are highlighted and annotated to show the previously known lineages (e.g., AFR1, AFR2), with the study samples distinguished from the representative reference genomes by the black circular tips (the reference genomes are tip-less, each colour denoting a unique *V. cholerae* O1 lineage. Where they occur, the distribution of study isolates found in each lineage is displayed next to the lineage name. Distinct clonal expansions with AFR12 and AFR15 arising from this study are denoted in red.

### Prevalence of acquired resistance genes and resistance mutations

The prevalence of genes encoding resistance to various antibiotics among the *Vibrio cholerae* study isolates is depicted in **Figure 4** and **Figure 5**. Trimethoprim resistance genes exhibited the highest prevalence, with 64 out of the 67 isolates (96%) harbouring at least one of four variants of the *dfrA* gene (**Figure 4** and **Figure 5**). *dfr*A1 occurred predominantly in the clinical isolates (40/46, 93%) (**Figure 4**). The environmental *V. cholerae* isolates, on the other hand, mainly harboured *dfr*A31 (7/24, 29%), along with *dfr*A15 (4/24, 17%) and *dfr*A1 (2/24, 8%) (**Figure 5**).

**Figure 4.**
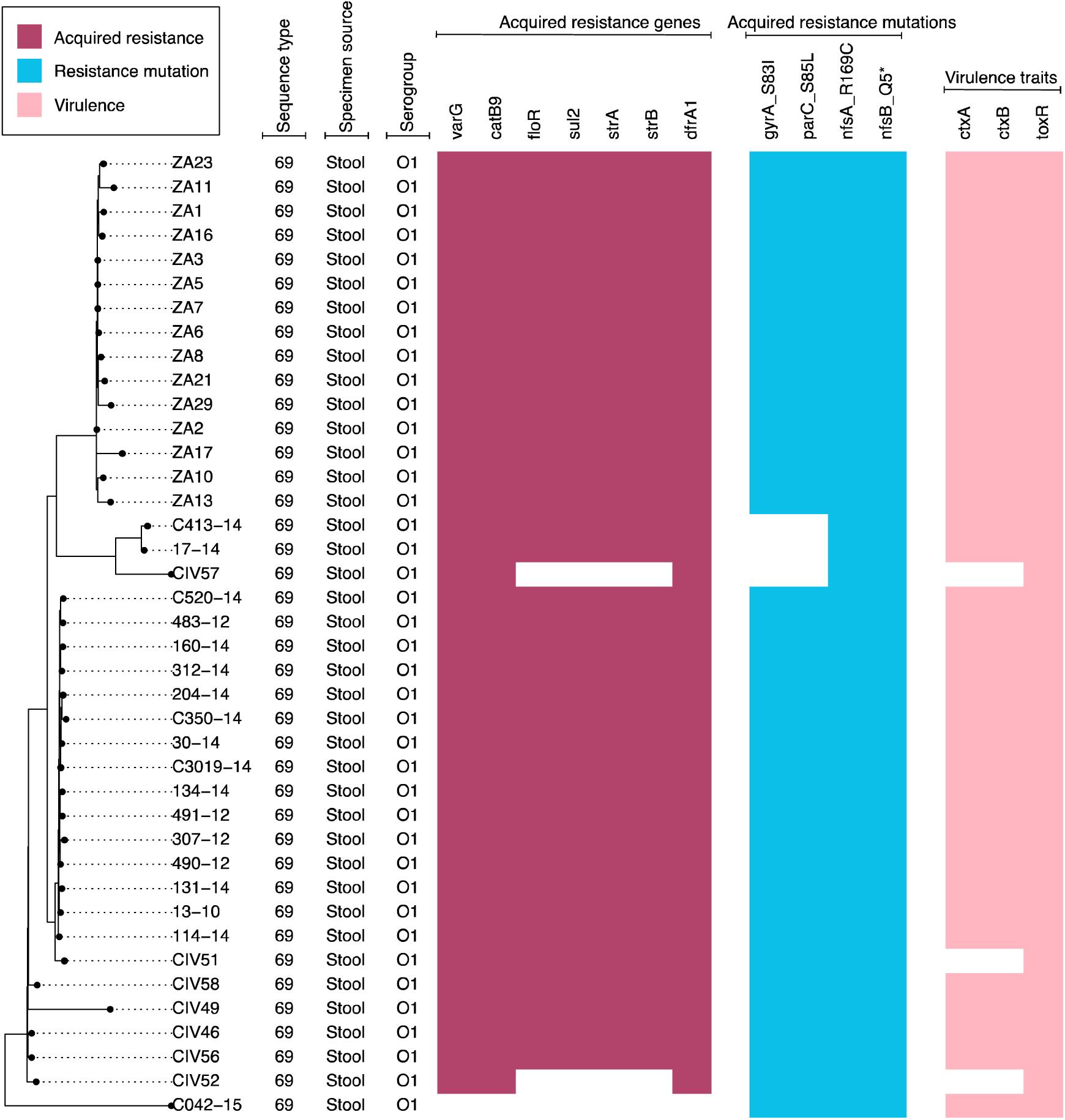
A neighbour-joining phylogenetic tree of the study *V. cholerae* ST69 serogroup O1 isolates (n=40) with antimicrobial resistance genes and virulence annotations. The phylogenetic tree depicts the evolutionary relationships among the study isolates, as determined by phylogenetic inference. The tree was reconstructed via Pathogenwatch (Reference #24), based on a concatenated alignment of 2,736 genes (3,075,173 bp) representing the core gene library for *V. cholerae* in Pathogenwatch. Each tip of the tree denotes a unique isolate, coloured by the geographic origin or source of the isolate (displayed in the legend). The heatmap visualises the presence of acquired antimicrobial resistance genes (raspberry), resistance mutations (blue raspberry) and virulence factors (light pink). The resistance genes, grouped by their classes, are as follows: carbapenems (*var*G), chloramphenicol (*cat*B9, *flo*R), aminoglycosides (*str*A, *str*B) and trimethoprim (*dfr*A1 and *dfr*A_new). Resistance mutations were as follows: quinolone resistance mutations (*gyr*A_S83I and *par*C_S85L) and nitrofuran (*nfs*A_R169C and *nfs*B_Q5*).

**Figure 5.**
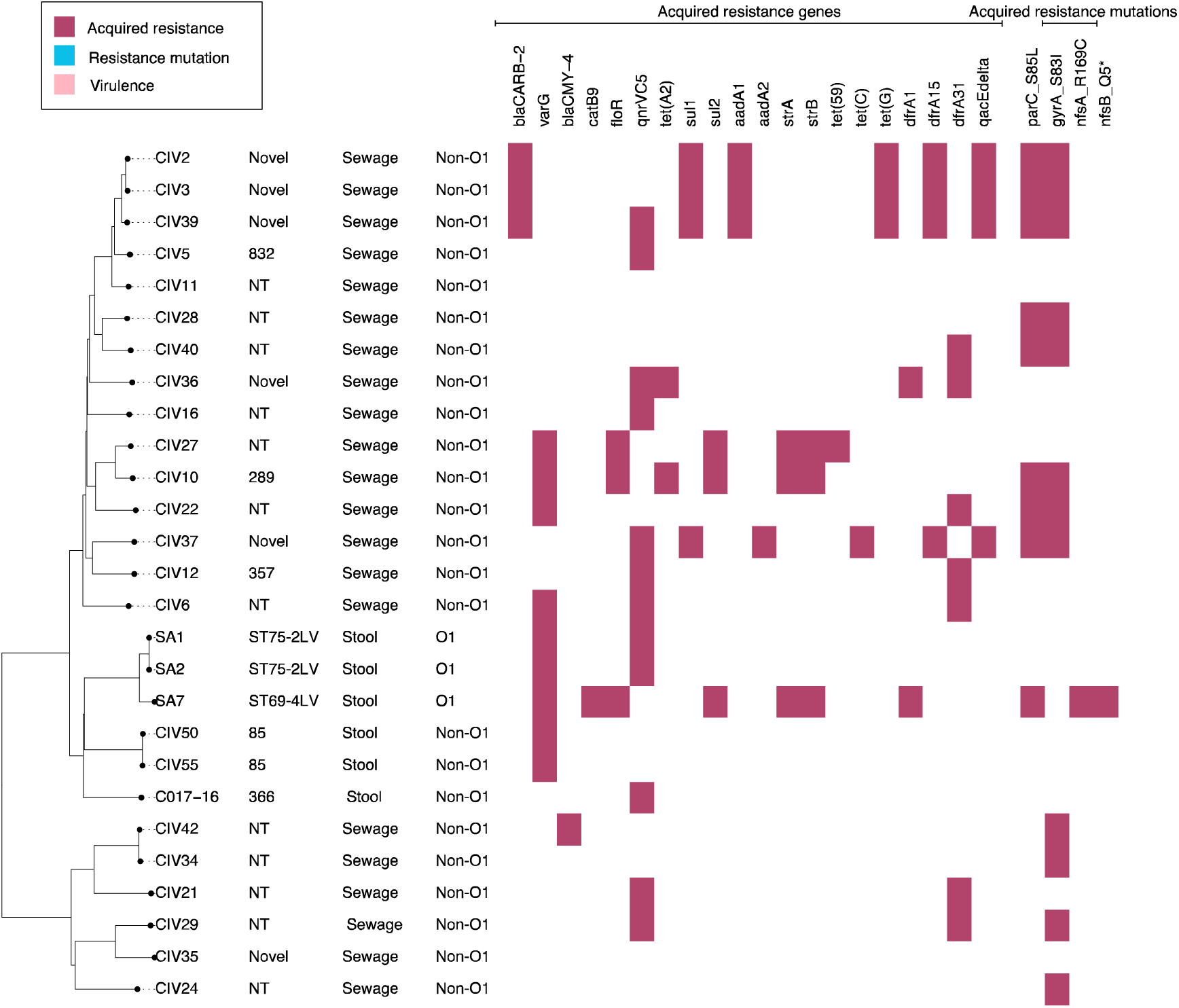
A neighbour-joining phylogenetic tree of the study *V. cholerae* non-ST69 isolates (n=27) with antimicrobial resistance genes and virulence annotations. The phylogenetic tree depicts the evolutionary relationships among the study isolates, as determined by phylogenetic inference. The tree was reconstructed via Pathogenwatch (Reference #24), based on a concatenated alignment of 2,736 genes (3,075,173 bp) representing the core gene library for *V. cholerae* in Pathogenwatch. Each tip of the tree denotes a unique isolate, coloured by the geographic origin or source of the isolate (displayed in the legend). The heatmap visualises the presence of acquired antimicrobial resistance genes (raspberry) and resistance mutations (blue raspberry). The resistance genes, grouped by their classes, are as follows: penicillins (*bl_a_*_CARB-2_), carbapenems (*var*G), third-generation cephalosporins (*bla*_CMY-4_), chloramphenicol (*cat*B9, *flo*R), sulphonamides (*sul*1*, Sul*2), aminoglycosides (*aad*A1, *aad*A2, *str*A, *str*B), tetracyclines (*tet*(C), *tet*(G), *tet*(59)), trimethoprim (*dfr*A1, *dfr*A15, *dfr*A31), and antiseptic (*qa*CEdelta). Resistance mutations were detected for quinolones as follows: (*gyr*A_S83I and *par*C_S85L

Conversely, some clinically relevant antibiotics demonstrated little or no resistance determinants among the study isolates. Notably, no determinants associated with azithromycin or rifampicin resistance were observed. Low prevalence rates were also seen for the antiseptic resistance gene *qac*Edelta, which occurred exclusively among the environmental strains (4/24, 16% of isolates). Similarly, tetracycline resistance genes occurred only among the environmental isolates, echoing findings from Algeria, Central African Republic, Kenya and Malawi, indicating widespread susceptibility of *V. cholerae* O1 isolates to tetracyclines (38–41). The genotypes included *tet*(C) in 4% of the environmental isolates (1/24), *tet*(G) in 13% (3/24), and *tet*(59) in 4% (1/24). The ampicillin resistance gene, *bla*_CARB-2_ gene, was found in the sewage isolates only (4/24, 12%).

The preponderance of the resistance determinants and mutations occurred among the *V. cholerae* O1 isolates (**Figure 4**), consistent with findings from other studies (38–42). For example, resistance determinants for chloramphenicol (*cat*B9 and *flo*R), aminoglycosides (*str*A and *str*B) and sulfamethoxazole (*sul*2) occurred in ≥88% (38/43) of the O1 strains but were detected in 4-16% of the non-O1 isolates (*cat*B9, 1/24, 4%; *flo*R, 3/24, 12%; *str*A, 3/24, 12%; *str*B, 2/24, 8%; *sul*1, 4/24, 16%; respectively). Conversely, the encoding resistance to ampicillin was identified in environmental isolates only (3/24, 12%), as was the *bla*_CMY-4_ gene encoding resistance to third-generation cephalosporins, which occurred in a single environmental isolate (1/24, 4%) (**Figure 5**).

Besides these, the *var*G resistance gene, a putative Ambler class B metallo-beta-lactamase found on the antibiotic resistance *var* regulon in *V. cholerae*, along with an antibiotic efflux pump (43), was present in 98% of the clinical *V. choloerae* O1 isolates (42/43) and 29% (7/24) of the environmental (non-O1) strains. However, we could not find much evidence for phenotypic carbapenem resistance due to *var*G in the literature; thus, further investigation is warranted to confirm this gene as a functional carbapenemase (43–45).

In addition to resistance genes, specific mutations conferring resistance to other antibiotic classes were identified. The *gyr*A_S83I and *par*C_S85L mutations, associated with quinolone resistance, were detected in 88% (38/43) of the serogroup O1 isolates. These isolates belonged to the current pandemic (7PET) clade and were distributed across the AFR12 and AFR15 lineages, consistent with the fixation of this trait in strains from these lineages (42). By contrast, the remaining serogroup O1 isolates, which lacked these fluoroquinolone resistance mutations, comprised three ST69 strains from the AFR9 lineage and two ST75-2LV clones from the pre-7PET clade.

Among non-O1 isolates, the *gyr*A_S83I and *par*C_S85L mutations were present in 50% (12/24) and 33% (8/24) of isolates, respectively. Resistance to furazolidone was mediated by the *nfs*A_R169C and *nfs*B_Q5* mutations, both of which were detected exclusively in the O1 isolates (41/43, 95%, all belonging to ST69) (**Figure 4** and **Figure 5**), indicating the fixation of these mutations in the O1 ST69 population.

Nearly all the *V. cholerae* O1 isolates (41/43, 95%) harboured resistance genes in at least three antibiotic classes, excluding *var*G (carbapenems). The high levels of resistance to critical antibiotics, particularly ciprofloxacin and nalidixic acid, raise significant concerns for treatment efficacy. However, the absence of resistance to tetracycline, azithromycin, and rifampicin among the V. cholerae O1 isolates suggests these antibiotics may remain effective treatment options.

Our findings mirrored a high prevalence of quinolone and nitrofuran resistance mutations and several resistance genes in the global population of *V. cholerae* O1 isolates (**File S4**). An analysis of n=4,858 *V. cholerae* O1 genomes hosted in Pathogenwatch (24) indicates that mutations *nfsB_Q5* and *nfsA_R169C*, conferring resistance to nitrofurans, are present in 70% of O1 isolates globally (3419/4858 and 3408/4858, respectively). Similarly, the *gyrA_S83I* and *parC_S85L* mutations occur in 63.96% (3107/4858) and 56.69% (2754/4858) of global isolates, respectively, reflecting the prevalence observed in our study where *gyrA* and *parC* mutations were present in 88% (38/43) of O1 isolates.

Our findings are consistent with the global prevalence of resistance genes, with *var*G detected in 99.77% (4847/4858) of all O1 strains, supporting the hypothesis that this gene is intrinsic to the *V. cholerae* population. Other notable resistance genes, such as *dfr*A1, *cat*B9, and *sul*2, encoding resistance to trimethoprim, chloramphenicol, and sulfamethoxazole, respectively, also exhibited similar global frequencies (*dfr*A1: 65.87% or 3200/4858; *cat*B9: 99.61% or 4839/4858; *sul*2: 63.59% or 3089/4858).

These findings highlight the widespread distribution of resistance traits in the global *V. cholerae* O1 population, likely driven by selective pressures and the dissemination of resistant lineages, particularly in clinical settings.

### Prevalence of virulence traits

Several virulence genes appeared to be conserved across the study isolates (**Figure 6**). These included the adhesion-associated genes, *rpo*S, *omp*U and the MSHA pilus; the haemagglutinin gene *hap*A, known for its cytotoxic and mucinolytic activities (46); the pore-forming toxin, *hly*A (haemolysin) (47); and *tox*R, the transmembrane transcription factor that regulates the production of virulence factors in *V. cholerae* (48), indicative that they were core to the species.

**Figure 6.**
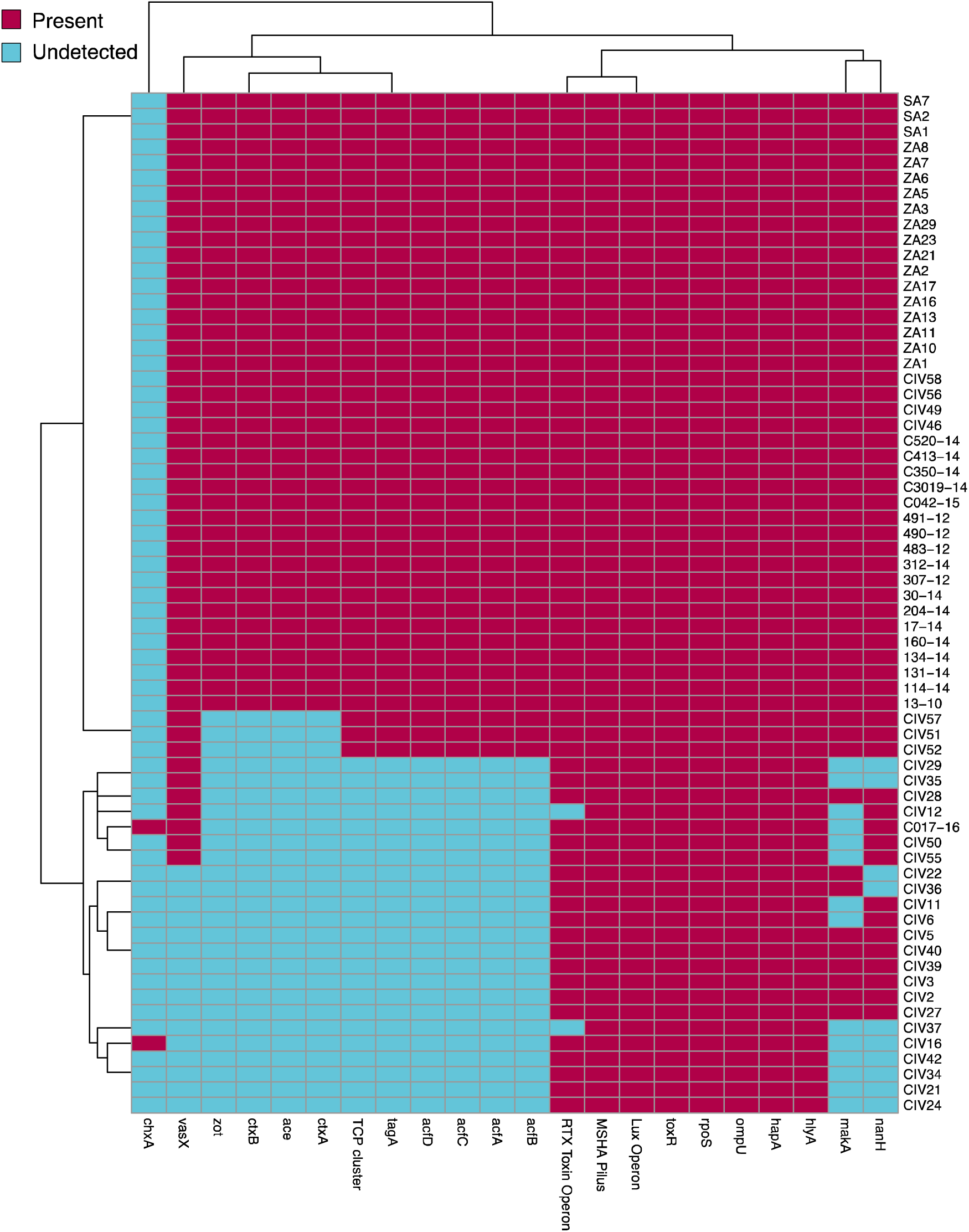
Binary Heatmap showing the presence and absence of virulence traits among the study *V. cholerae* isolates. The heatmap displays the presence (raspberry) and absence (soft cyan) of specific virulence genes across the study isolates. Each row represents a sample, and each column represents a gene of interest, as identified via Pathogenwatch and TheiaProk. The resistance genes, by their classes, are as follows: *rpo*S, a stress gene involved in adhesion; *omp*U, an outer membrane protein also involved in adhesion; MSHA Pilus, a pili structure that promotes adhesion to host surfaces; *<u>hap</u>*A, a haemagglutinin/protease gene that facilitates bacterial invasion and immune evasion; *hly*A, a haemolysin gene that contributes to cytotoxicity; *tox*R, a transmembrane transcriptional activator that regulates the production of virulence factors, including cholera toxin; *ctx*A and *ctx*B, the cholera toxin genes responsible for cholera toxin production; ace, a toxin gene associated with cholera enterotoxin; *acf*A, *acf*B, *acf*C, and *acf*D, colonisation factor genes necessary for intestinal colonisation; *zot*, a toxin gene that increases intestinal permeability; *chx*A, a toxin gene; and *vas*X, a gene encoding a toxin involved in the type VI secretion system, aiding bacterial virulence.

Besides these, several virulence traits appeared to be fixed among the clinical *V. cholerae* O1 study isolates. For example, the cholera toxin genes *ctx*A and *ctx*B, which produce the ‘rice-water’ stools characteristic of cholera (49, 50); the toxin gene *ace*; the colonisation operon *acf*A*, acf*B, *acf*C, *acf*D; the toxin co-regulated pilus (TCP) cluster, associated with colonisation in humans and animals (51), and the toxin gene *zot* occurred in >93 % (40/43) of the *V. cholerae* O1 isolates but were not detected among any of the non-O1 isolates. By contrast, the cholix toxin gene, *chx*A (an exotoxin) (52), was found in 8% (2/24) of the environmental isolates but was not found among any of the clinical isolates. A previous study found *chx*A in both O1 (clinical) and non-O1 (environmental) strains (53). The complete list of virulence traits detected among the study *Vibrio* isolates is provided in **File S5**.

## Discussion

This study demonstrates the utility of Oxford Nanopore Technology (ONT) sequencing for enhancing genomic surveillance of *V. cholerae* in Africa, a critical capability for networks like PulseNet Africa. By applying ONT sequencing, we were able to generate high-resolution data on the genetic diversity, AMR profiles, and virulence traits of *V. cholerae* isolates from multiple African countries. This approach has significant potential for strengthening regional surveillance efforts for foodborne pathogens, including *V. cholerae*, as well as other critical pathogens under PulseNet Africa’s remit. The ability to rapidly generate and analyse whole-genome sequences using ONT in resource-limited settings could enhance the early detection of outbreaks, track the spread of AMR, and improve public health responses across the continent.

Among the clinical isolates analysed, three non-O1 strains were retrieved from cases presenting with cholera-like symptoms. In addition, several study isolates were misidentified as *V. cholerae*, including non-*Vibrio* species such as *K. quasipneumoniae*, *E. hormaechei*, and *A. enteropelogenes*. These findings underscore a significant diagnostic challenge: pathogens causing cholera-like disease are frequently misidentified as *V. cholerae* by conventional microbiological methods. Previous studies in Africa and elsewhere have suggested that certain enteric bacteria can cause cholera-like diarrheal infections, leading to their misattribution to *V. cholerae* during cholera outbreaks (54–57). This issue is further underscored by findings from a large cholera outbreak in Malawi, where Chaguza et al. (2024) identified non-*V. cholerae* species in approximately 28% of cases (19 out of 68) (39).

The identification of clinically relevant *Vibrio* species, such as *V. fluvialis*, *V. furnissii*, and *V. navarrensis*—all of which cause gastrointestinal illness (58)—is particularly noteworthy in the context of horizontal gene transfer among *Vibrio* species (59). For instance, *V. mimicus*, though a non-*cholerae* species, can harbour the cholera toxin (*ctx*) gene (60), highlighting the potential for genetic exchange that may influence virulence. Similarly, *V. fluvialis* has been shown to possess an El Tor-like hemolysin (*hly*A), a gene associated with pathogenicity (61), underscoring the importance of monitoring these species for genetic traits relevant to outbreaks and public health.

From this study, the *V. furnissii* isolate was recovered from a clinical stool specimen, while *V. fluvialis* and *V. navarrensis* isolates were all sourced from sewage samples. These findings emphasise the necessity of accurate genomic characterisation and expanded surveillance for *Vibrio* species, especially given the challenges of speciation due to phenotypic heterogeneity even within a single *Vibrio* species. Genomic surveillance will be critical to improving pathogen detection, understanding their evolutionary dynamics, and mitigating risks posed by emerging strains.

An earlier survey of global *V. cholerae* diversity by Weill et al. (2017) (11) included strains from Côte d’Ivoire (n=29, spanning 1970–2006), Zambia (n=26, spanning 1996–2012), and Ghana (n=8, spanning 1970–2014). Our study extends this timeline, including isolates collected between 2010 and 2024, and provides new insights into the genomic diversity of *V. cholerae* O1 in these regions.

The phylogenetic analysis uncovered significant genetic diversity among the isolates, with several forming novel clades, particularly within the *V. cholerae* O1 serogroup. Most clinical isolates clustered within the pandemic 7PET lineage, consistent with previous studies on *V. cholerae* transmission in Africa (4, 8, 9, 11, 15, 41). However, the discovery of distinct novel sublineages among clinical O1 isolates (35/45, 77.8%) underscores the potential for ongoing, uncharacterised transmission events or the importation of new strains into the region. Consistent with recent reports from South Africa, the two ST75-2LV strains from our study clustered closely with isolates from the pre-7PET clade. The closest relative to these strains was the US Gulf Coast *V. cholerae* O1 clone (37), highlighting the continued dominance of ST75 in cholera cases reported in South Africa. The environmental strains also displayed considerable diversity, with the 24 isolates being spread over five distinct STs and six potentially novel STs. Such diversity underscores the importance of continuously monitoring these populations to detect emergent strains that may acquire virulence factors or resistance genes.

The clustering of non-O1 isolates into separate groups, particularly those recovered from sewage samples, indicates a divergence in environmental strains from those driving outbreaks. At least in our study, environmental strains do not appear to seed outbreaks, as they are genetically quite divergent from the clinical strains, although they harbour some key virulence traits. Further investigation into the role of environmental reservoirs as potential contributors to future outbreaks and their broader role in cholera epidemiology is warranted (62–65).

The high prevalence of AMR genes in the study isolates is concerning, especially given the widespread presence of resistance determinants for clinically essential antibiotics such as trimethoprim, ciprofloxacin, and nalidixic acid. The near-universal presence of *dfrA* alleles and high prevalence of quinolone resistance mutations (*gyr*A*_*S83I and *par*C*_*S85L) suggest that these drugs may be increasingly ineffective for cholera treatment in the region. This finding mirrors global trends of rising resistance to quinolones and other frontline antibiotics (4, 11, 66), which may complicate treatment strategies and necessitate a shift in therapeutic approaches.

The detection of AMR genes in all O1 isolates, with each exhibiting resistance to at least three classes of antibiotics, is consistent with findings from other African studies (38–42) and the global *V. cholerae* O1 populations and raises concerns about the continued efficacy of current treatment protocols. While no resistance determinants were identified for azithromycin and rifampicin, the growing resistance to other antibiotics used in cholera treatment, such as tetracyclines and quinolones, suggests that alternative strategies must be developed. The absence of azithromycin resistance among these isolates offers a potential avenue for maintaining treatment efficacy, as this drug may remain a viable option for first-line treatment (4, 6).

The stark contrast in resistance profiles between O1 and non-O1 isolates further emphasises the complexity of cholera epidemiology in Africa. O1 isolates exhibited significantly higher levels of AMR, with many harbouring resistance determinants to multiple classes of antibiotics. This difference may reflect the clinical settings from which O1 isolates were derived, as these strains are more likely to have been exposed to selective pressures from antibiotic use in humans. In contrast, non-O1 isolates, particularly those from environmental sources, displayed comparatively lower levels of resistance, which could suggest reduced exposure to antibiotics in these reservoirs. However, the potential for environmental strains to acquire resistance genes through horizontal gene transfer cannot be overlooked and requires ongoing monitoring. While the environmental strains displayed fewer resistance genes compared to clinical isolates, there were overlaps in certain determinants; e.g., the quinolone mutations, *gyr*A*_*S83I and *par*C*_*S85L, occurred in 50% and 33% of the environmental strains, respectively. This suggests some shared AMR determinants between the two groups, which is crucial to further investigate the direction of AMR transmission within the environment and the potential role of environmental strains in the spread of resistance.

Although more virulence genes were found in the clinical strains, with several being fixed in the population but absent from the environmental strains, there were key overlaps in virulence genes among both O1 and non-O1 strains, warranting continuous monitoring. The high prevalence of virulence genes among the O1 isolates confirms their pathogenic potential and role in driving cholera outbreaks in the region.

## Limitations

A major limitation of this study is that the sample collection relied on available archived and viable isolates from four countries. As a result, the collection may not fully represent the diversity of circulating *Vibrio cholerae* strains in the region during the study period. The lack of phenotypic antimicrobial susceptibility test data is another limitation. Although we were able to analyse the genotypic profiles of AMR, phenotypic data would have provided direct evidence of resistance patterns in the isolates. Unfortunately, logistic constraints did not allow us to conduct phenotypic testing, which limits the study’s ability to fully assess the clinical implications of the detected resistance genes.

## Conclusion

Our study demonstrates the value of using WGS to uncover the genomic diversity, AMR profiles, and virulence traits of *V. cholerae* strains in Africa. Further work is needed to develop standardised QC metrics and standard operating procedures for handling ONT sequencing data within PulseNet Africa and its sister networks. Establishing these protocols will ensure data consistency, reliability, and comparability across labs, thereby strengthening the regional genomic surveillance framework for *Vibrio cholerae* and other pathogens.

The identification of clonal expansion within certain lineages, characterised by high levels of AMR genotypes in clinical isolates, provides essential information for improving cholera surveillance and response strategies. Given the high prevalence of resistance to critical antibiotics such as trimethoprim and quinolones and the potential for further AMR evolution, it is imperative to reassess treatment guidelines and enhance genomic monitoring.

## Acknowledgements

The authors express our heartfelt gratitude to the University Teaching Hospitals (UTH) management, the Zambia National Public Health Institute (ZNPHI), and the National Health Research Authority (NHRA) Zambia for granting permission to utilise DNA for this collaborative work. We sincerely thank the Institut Pasteur of Côte d’Ivoire, the District of Abidjan, Dr. Yao Kouamé Eric, and Mr Koffi Romain from the Environmental Chemistry and Microbiology Unit, National Institute of Public Hygiene, for their invaluable contributions to this study. Thanks also go to all the laboratory scientists and staff of the National Public Health Laborotory, Korle-Bu, Accra.

We are also deeply appreciative of the support provided by the Unit Leadership Board and the Laboratory Management and Training Department of MRCG at LSHTM for hosting the training workshop, as well as the dedicated support staff. Special recognition goes to the research support staff, the health and safety team, the finance team, travel officer Fanta Singhateh, and the Project Managers Nfamara Camara and Landing Ndow for their exceptional assistance and dedication to the success of this work.

Finally, we would like to acknowledge Angela Poates (APHL), Morgan Schroeder and Kristy Kubota (CDC), Michele Scribner, Emma Doughty, and James Otieno (Theiagen Genomics) for their invaluable contributions to the planning and execution of the training workshop.

## Author contributions

Conceptualization – Initial idea, hypothesis development, and overall research design: EFN, PS, JR, HC.

Data Collection – Conducting experiments, collecting samples, gathering or analysing data: SAB, NEA, RA, JCBA, LA, YB, GB, DC, RC, KJC, FAD, DD, CD, MD, MMD, SF, RJ, JBK, AK, HFK, DK, NKNK, CL, HM, GM, JM, FN, GOO, JR, AS, AKS, AS, PS, DS, NT, PLST, PEMT, CS, SNV, DV.

Methodology – Development or design of the methodology; selecting and designing the research methods: EFN, PS, JR.

Writing - Original Draft – Writing the initial draft of the manuscript: EFN.

Writing - Review & Editing – Reviewing, editing, and revising the manuscript: EFN, SAB, NEA, RA, JCBA, LA, YB, GB, HC, DC, RC, KJC, FAD, DD, CD, MD, MMD, SF, KEH, RJ, JBK, AK, HFK, DK, NKNK, CL, HM, GM, JM, FN, GOO, JR, AS, AKS, AS, PS, DS, NT, PLST, PEMT, CS, SNV, DV.

## Ethical statement

The inclusion of country-specific data in this study was approved by the institutional review boards and respective ministries of health in the four countries that contributed samples. Informed patient consent was waived as the samples were collected as part of routine diagnostic procedures. All patient data associated with these isolates were anonymised to ensure no possibility of patient identification based on age, sex, or hospital-related information. This research was conducted in accordance with the principles outlined in the Declaration of Helsinki.

---

**Figure S1:**
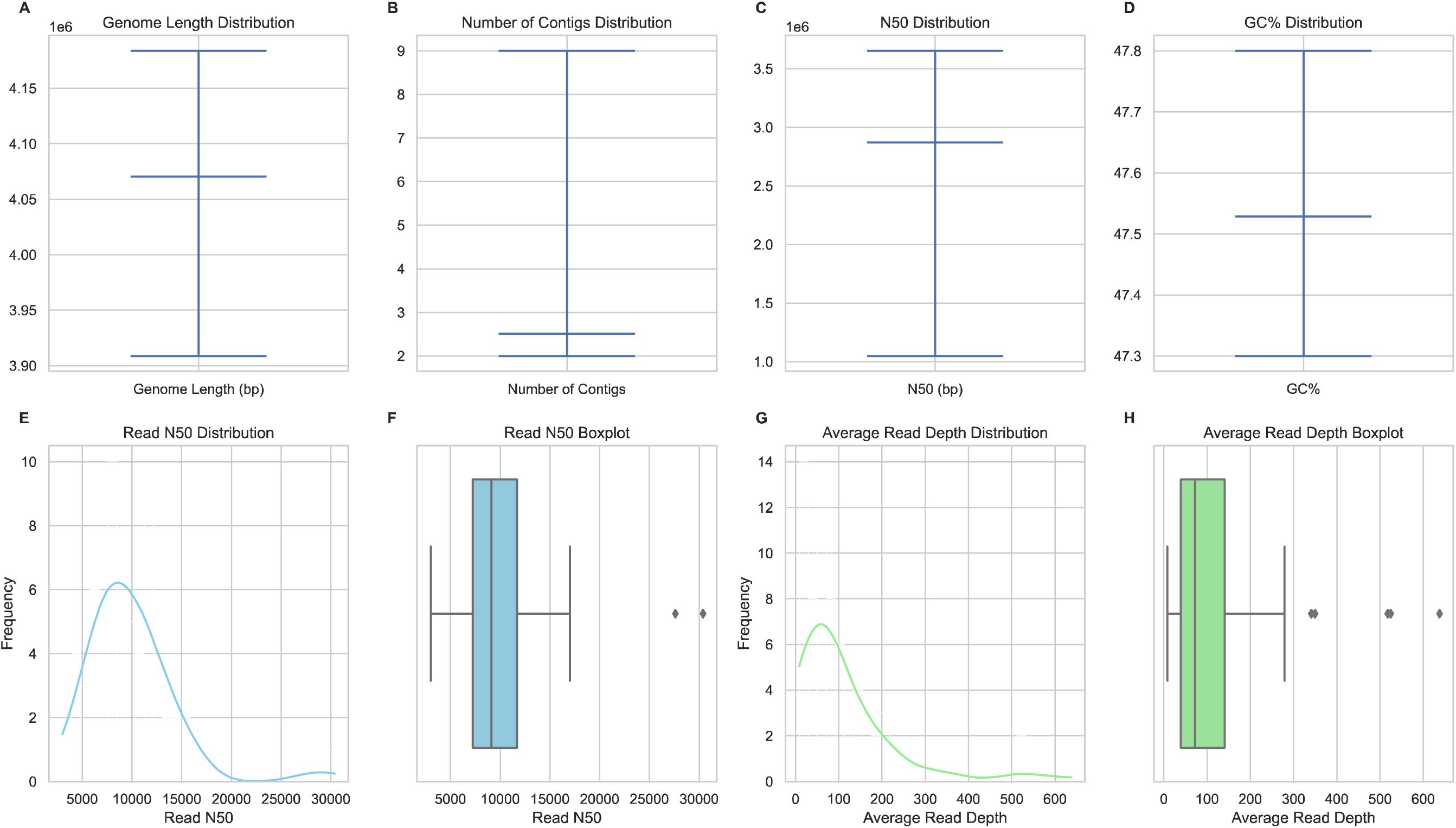
Distribution of Read and Genome Assembly Metrics Panels A–D: Violin plots showing the distribution of genome assembly metrics across *Vibrio cholerae* samples (n=67), as follows: **A.** Genome length distribution (in base pairs). **B.** Number of contigs distribution. **C.** N50 distribution**. D.** GC% distribution across genome assemblies**. Panels E–H:** Read quality and coverage metrics for ONT sequencing: **E.** Histogram showing the distribution of read N50 values. **F.** Read N50 boxplot summarising the range and outliers in read N50 values. **G.** Histogram depicting the average read depth distribution. **H.** Boxplot illustrating the spread and outliers in average read depth values.

**Figure S2.**
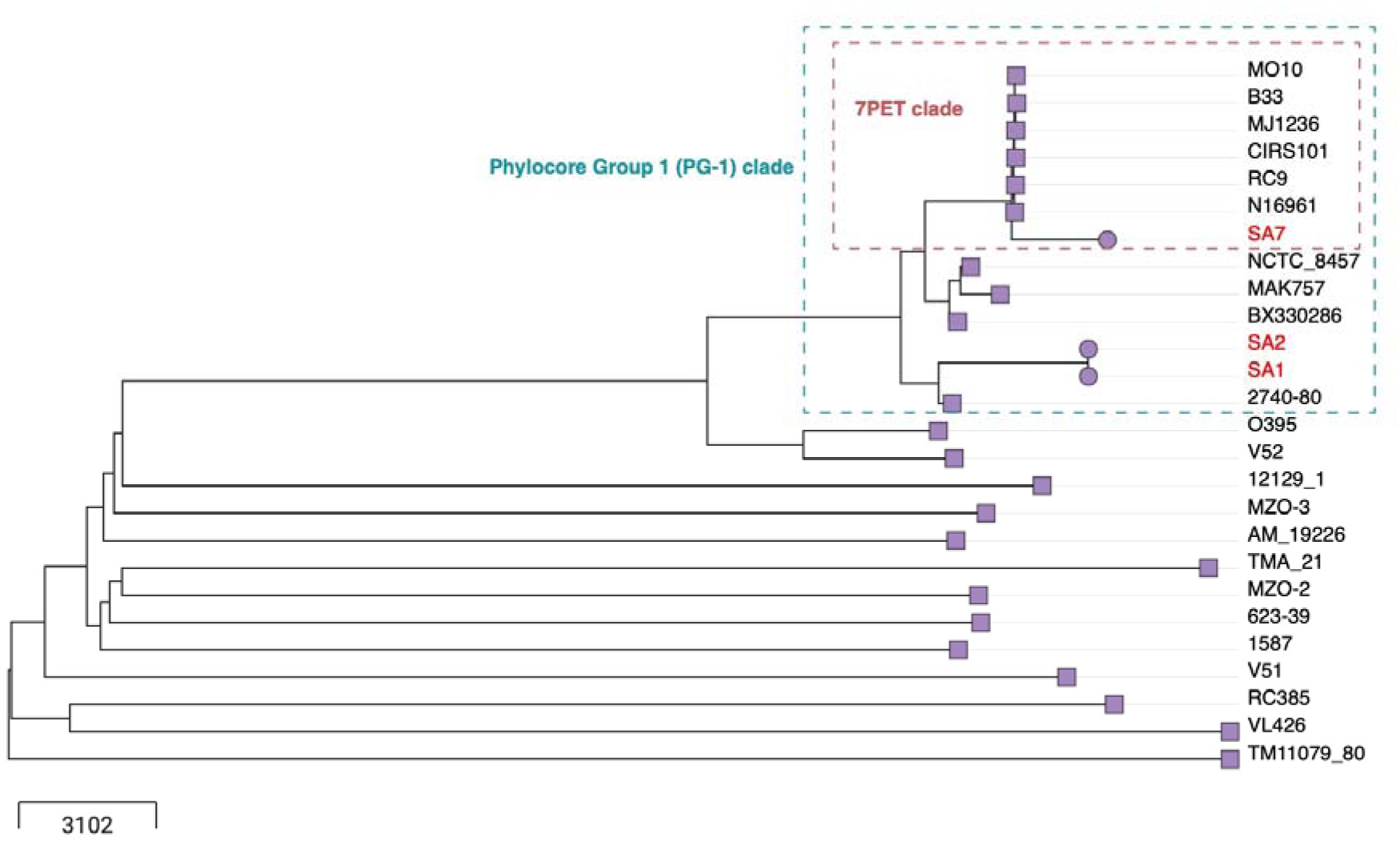
A Neighbour-Joining tree illustrating the phylogenetic placement of the study isolates ST-69-4LV and ST-75-2LV. The tree was reconstructed using Vibriowatch (Reference #24) and incorporates the collection of isolates published by Chun et al. (Reference #29). Study isolates are represented by circular tips with red labels, while reference isolates from the Chun collection are represented by square tips and black labels. The soft brick red dashed rectangle highlights isolates belonging to the current global “7th pandemic” (7PET) clade, whereas the teal rectangle marks the broader Phylocore Group 1 (PG- 1) clade. Notably, the two ST-75-2LV study isolates cluster closely with NCTC_8457, MAK757, and BX330286, which are sometimes referred to as ‘pre-7PET’ isolates.

**Figure S3:**
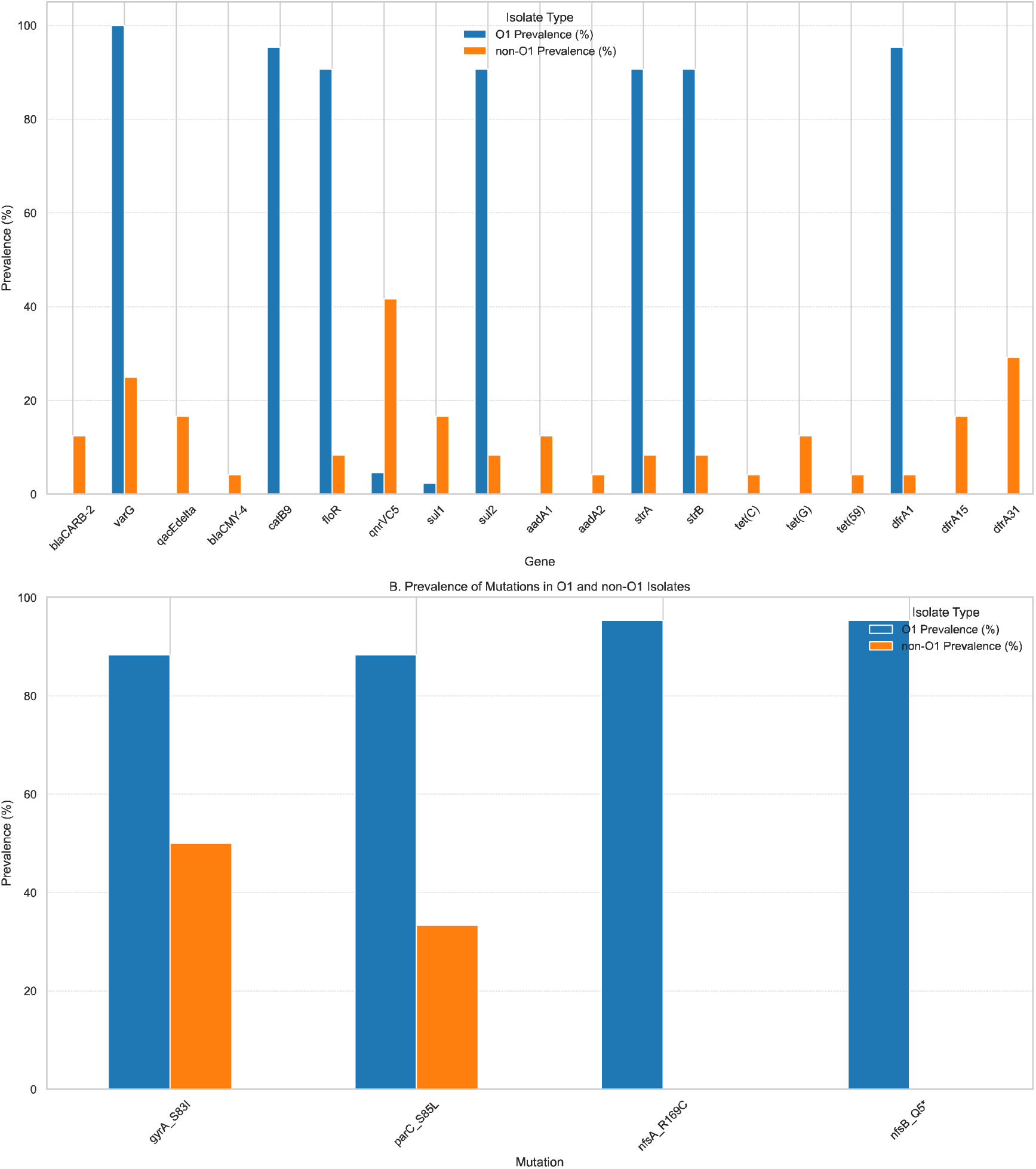
Prevalence of resistance genes and mutations in O1 vs non-O1 isolates. (**A**) The bar plot shows the prevalence of resistance genes across O1 and non-O1 isolates. The blue bars represent the prevalence of resistance genes in O1 isolates, while the orange bars represent the prevalence in non-O1 isolates. The genes are grouped by antibiotic resistance classes, including penicillins (*blaCARB-2*), carbapenems (*var*G), third-generation cephalosporins (*bla*_CMY-4_), chloramphenicol (*cat*B9, *flo*R), aminoglycosides (*aad*A1, *aad*A2, *str*A, *str*B), tetracyclines (*tet*(C), *tet*(G), *tet*(59), and trimethoprim (*dfr*A1, *dfr*A15, dfrA31, *dfr*A_new). (B) The bar plot shows the prevalence of key mutations associated with antibiotic resistance in O1 and non-O1 isolates. The blue bars represent the prevalence in O1 isolates, and the orange bars represent non-O1 isolates. The mutations include those associated with quinolone resistance (*gyr*A_S83I, *par*C_S85L) and furazolidone resistance (*nfs*A_R169C, *nfs*B_Q5*). Data labels indicate the precise prevalence percentages.

## References

1. Mwaba J, Ferreras E, Chizema-Kawesa E, Mwimbe D, Tafirenyika F, Rauzier J, et al. Evaluation of the SD bioline cholera rapid diagnostic test during the 2016 cholera outbreak in Lusaka, Zambia. Trop Med Int Health. 2018;23(8):834–40.

2. Mengel MA, Delrieu I, Heyerdahl L, Gessner BD. Cholera outbreaks in Africa. Curr Top Microbiol Immunol. 2014;379:117–44.

3. Kanungo S, Azman AS, Ramamurthy T, Deen J, Dutta S. Cholera. Lancet. 2022;399(10333):1429–40.

4. Mashe T, Domman D, Tarupiwa A, Manangazira P, Phiri I, Masunda K, et al. Highly resistant cholera outbreak strain in Zimbabwe. N Engl J Med. 2020;383(7):687–9.

5. Mohanraj RS, Samanta P, Mukhopadhyay AK, Mandal J. Haitian-like genetic traits with creeping MIC of Azithromycin in *Vibrio cholerae* O1 isolates from Puducherry, India. J Med Microbiol. 2020;69(3):372–8.

6. Parvin I, Shahunja KM, Khan SH, Alam T, Shahrin L, Ackhter MM, et al. Changing susceptibility pattern of *Vibrio cholerae* O1 isolates to commonly used antibiotics in the largest diarrheal disease hospital in Bangladesh during 2000-2018. Am J Trop Med Hyg. 2020;103(2):652–8.

7. Colwell RR. Global climate and infectious disease: the cholera paradigm. Science. 1996;274(5295):2025-31.

8. Smith AM, Sekwadi P, Erasmus LK, Lee CC, Stroika SG, Ndzabandzaba S, et al. Imported cholera cases, South Africa, 2023. Emerg Infect Dis. 2023;29(8):1687-90.

9. Xiao S, Abade A, Boru W, Kasambara W, Mwaba J, Ongole F, et al. New *Vibrio cholerae* sequences from Eastern and Southern Africa alter our understanding of regional cholera transmission. medRxiv. 2024.

10. Ekeng E, Tchatchouang S, Akenji B, Issaka BB, Akintayo I, Chukwu C, et al. Regional sequencing collaboration reveals persistence of the T12 *Vibrio cholerae* O1 lineage in West Africa. Elife. 2021;10.

11. Weill FX, Domman D, Njamkepo E, Tarr C, Rauzier J, Fawal N, et al. Genomic history of the seventh pandemic of cholera in Africa. Science. 2017;358(6364):785-9.

12. World Health Organization. Whole genome sequencing as a tool to strengthen foodborne disease surveillance and response: module 1: introductory module. 2023.

13. World Health Organization. GLASS whole-genome sequencing for surveillance of antimicrobial resistance. 2020.

14. World Health Organization. Whole genome sequencing for foodborne disease surveillance: Landscape paper. 2018.

15. Mwaba J, Debes AK, Murt KN, Shea P, Simuyandi M, Laban N, et al. Three transmission events of *Vibrio cholerae* O1 into Lusaka, Zambia. BMC Infect Dis. 2021;21(1):570.

16. De R, Ghosh JB, Sen Gupta S, Takeda Y, Nair GB. The role of *Vibrio cholerae* genotyping in Africa. J Infect Dis. 2013;208 Suppl 1:S32–8.

17. Deng X, den Bakker HC, Hendriksen RS. Genomic epidemiology: Whole-genome-sequencing-powered surveillance and outbreak investigation of foodborne bacterial pathogens. Annu Rev Food Sci Technol. 2016;7:353–74.

18. Carleton HAG-S, P. Whole-genome sequencing is taking over foodborne disease surveillance. Microbe Magazine 2016. p. 311–7

19. Davedow T, Carleton H, Kubota K, Palm D, Schroeder M, Gerner-Smidt P, et al. PulseNet International survey on the implementation of whole genome sequencing in low and middle-income countries for foodborne disease surveillance. Foodborne Pathog Dis. 2022;19(5):332–40.

20. Hounmanou YMG, Leekitcharoenphon P, Kudirkiene E, Mdegela RH, Hendriksen RS, Olsen JE, et al. Genomic insights into *Vibrio cholerae* O1 responsible for cholera epidemics in Tanzania between 1993 and 2017. PLoS Negl Trop Dis. 2019;13(12):e0007934.

21. Domman D, Quilici ML, Dorman MJ, Njamkepo E, Mutreja A, Mather AE, et al. Integrated view of *Vibrio cholerae* in the Americas. Science. 2017;358(6364):789-93.

22. Hounmanou YMG, Leekitcharoenphon P, Hendriksen RS, Dougnon TV, Mdegela RH, Olsen JE, et al. Surveillance and genomics of toxigenic *Vibrio cholerae* O1 From fish, phytoplankton and water in Lake Victoria, Tanzania. Front Microbiol. 2019;10:901.

23. Connor TR, Loman NJ, Thompson S, Smith A, Southgate J, Poplawski R, et al. CLIMB (the Cloud Infrastructure for Microbial Bioinformatics): an online resource for the medical microbiology community. Microb Genom. 2016;2(9):e000086.

24. Vibriowatch, a Pathogenwatch database for *Vibrio cholerae*. Available from: https://vibriowatch.readthedocs.io/en/latest/index.html.

25. Wingett SW, Andrews S. FastQ Screen: A tool for multi-genome mapping and quality control. F1000Res. 2018;7:1338.

26. Bouras G, Houtak G, Wick RR, Mallawaarachchi V, Roach MJ, Papudeshi B, et al. Hybracter: enabling scalable, automated, complete and accurate bacterial genome assemblies. Microb Genom. 2024;10(5).

27. Lumpe J, Gumbleton L, Gorzalski A, Libuit K, Varghese V, Lloyd T, et al. GAMBIT (Genomic Approximation Method for Bacterial Identification and Tracking): A methodology to rapidly leverage whole genome sequencing of bacterial isolates for clinical identification. PLoS One. 2023;18(2):e0277575.

28. Seemann T. Abricate. Available from: https://github.com/tseemann/abricate.

29. Carattoli A, Zankari E, García-Fernández A, Voldby Larsen M, Lund O, Villa L, et al. In silico detection and typing of plasmids using PlasmidFinder and plasmid multilocus sequence typing. Antimicrob Agents Chemother. 2014;58(7):3895–903.

30. Lees JA, Harris SR, Tonkin-Hill G, Gladstone RA, Lo SW, Weiser JN, et al. Fast and flexible bacterial genomic epidemiology with PopPUNK. Genome Res. 2019;29(2):304–16.

31. Chun J, Grim CJ, Hasan NA, Lee JH, Choi SY, Haley BJ, et al. Comparative genomics reveals mechanism for short-term and long-term clonal transitions in pandemic *Vibrio cholerae*. Proc Natl Acad Sci U S A. 2009;106(36):15442–7.

32. Derelle R, von Wachsmann J, Mäklin T, Hellewell J, Russell T, Lalvani A, et al. Seamless, rapid and accurate analyses of outbreak genomic data using Split K-mer Analysis (SKA). bioRxiv. 2024:2024.03.25.586631.

33. Jamin C, De Koster S, van Koeveringe S, De Coninck D, Mensaert K, De Bruyne K, et al. Harmonization of whole-genome sequencing for outbreak surveillance of. Microb Genom. 2021;7(7).

34. Ashton P. Validation of SKA analysis for genomic epidemiology. 2021. Available from: https://bitsandbugs.org/2021/07/08/validation-of-ska-analysis-for-genomic-epidemiology/.

35. Stamatakis A. RAxML-VI-HPC: maximum likelihood-based phylogenetic analyses with thousands of taxa and mixed models. Bioinformatics. 2006;22(21):2688–90.

36. Pathogenwatch. Speciator. Available from https://pathogen.watch/.

37. Smith AM, Weill FX, Njamkepo E, Ngomane HM, Ramalwa N, Sekwadi P, et al. Emergence of *Vibrio cholerae* O1 sequence type 75, South Africa, 2018-2020. Emerg Infect Dis. 2021;27(11):2927–31.

38. Shah MM, Bundi M, Kathiiko C, Guyo S, Galata A, Miringu G, et al. Antibiotic-resistant *Vibrio cholerae* O1 and its SXT elements associated with two cholera epidemics in Kenya in 2007 to 2010 and 2015 to 2016. Microbiol Spectr. 2023;11(3):e0414022.

39. Chaguza C, Chibwe I, Chaima D, Musicha P, Ndeketa L, Kasambara W, et al. Genomic insights into the 2022-2023 Vibrio cholerae outbreak in Malawi. Nat Commun. 2024;15(1):6291.

40. Breurec S, Franck T, Njamkepo E, Mbecko JR, Rauzier J, Sanke-Waïgana H, et al. Seventh pandemic *Vibrio cholerae* O1 sublineages, Central African Republic. Emerg Infect Dis. 2021;27(1):262–6.

41. Benamrouche N, Belkader C, Njamkepo E, Zemam SS, Sadat S, Saighi K, et al. Outbreak of imported seventh pandemic *Vibrio cholerae* O1 El Tor, Algeria, 2018. Emerg Infect Dis. 2022;28(6):1241–5.

42. Rouard C, Greig DR, Tauhid T, Dupke S, Njamkepo E, Amato E, et al. Genomic analysis of *Vibrio cholerae* O1 isolates from cholera cases, Europe, 2022. Euro Surveill. 2024;29(36).

43. Lin HV, Massam-Wu T, Lin CP, Wang YA, Shen YC, Lu WJ, et al. The *Vibrio cholerae* var regulon encodes a metallo-β-lactamase and an antibiotic efflux pump, which are regulated by VarR, a LysR-type transcription factor. PLoS One. 2017;12(9):e0184255.

44. Goh JXH, Tan LT, Law JW, Khaw KY, Ab Mutalib NS, He YW, et al. Insights into carbapenem resistance in *Vibrio* species: Current status and future perspectives. Int J Mol Sci. 2022;23(20).

45. Monir MM, Islam MT, Mazumder R, Mondal D, Nahar KS, Sultana M, et al. Genomic attributes of *Vibrio cholerae* O1 responsible for 2022 massive cholera outbreak in Bangladesh. Nat Commun. 2023;14(1):1154.

46. Benitez JA, Silva AJ. *Vibrio cholerae* hemagglutinin(HA)/protease: An extracellular metalloprotease with multiple pathogenic activities. Toxicon. 2016;115:55–62.

47. Williams SG, Manning PA. Transcription of the *Vibrio cholerae* haemolysin gene, *hly*A, and cloning of a positive regulatory locus, hlyU. Mol Microbiol. 1991;5(8):2031–8.

48. Gubensäk N, Schrank E, Hartlmüller C, Göbl C, Falsone FS, Becker W, et al. Structural and DNA-binding properties of the cytoplasmic domain of *Vibrio cholerae* transcription factor ToxR. J Biol Chem. 2021;297(4):101167.

49. Harris JB, LaRocque RC, Qadri F, Ryan ET, Calderwood SB. Cholera. Lancet. 2012;379(9835):2466-76.

50. Gorbach SL, Banwell JG, Jacobs B, Chatterjee BD, Mitra R, Brigham KL, et al. Intestinal microflora in Asiatic cholera. I. “Rice-water” stool. J Infect Dis. 1970;121(1):32–7.

51. Manning PA. The tcp gene cluster of *Vibrio cholerae*. Gene. 1997;192(1):63–70.

52. Awasthi SP, Asakura M, Chowdhury N, Neogi SB, Hinenoya A, Golbar HM, et al. Novel cholix toxin variants, ADP-ribosylating toxins in *Vibrio cholerae* non-O1/non-O139 strains, and their pathogenicity. Infect Immun. 2013;81(2):531–41.

53. Purdy AE, Balch D, Lizárraga-Partida ML, Islam MS, Martinez-Urtaza J, Huq A, et al. Diversity and distribution of cholix toxin, a novel ADP-ribosylating factor from *Vibrio cholerae*. Environ Microbiol Rep. 2010;2(1):198–207.

54. Gurwith M, Bourque C, Cameron E, Forrest G, Green M. Cholera-like diarrhea in Canada. Report of a case associated with enterotoxigenic *Escherichia coli* and a toxin-producing *Aeromonas hydrophila*. Arch Intern Med. 1977;137(10):1461–4.

55. Mendes-Marques CL, Nascimento LM, Theophilo GN, Hofer E, Melo Neto OP, Leal NC. Molecular characterization of *Aeromonas* spp. and *Vibrio cholerae* O1 isolated during a diarrhea outbreak. Rev Inst Med Trop Sao Paulo. 2012;54(6):299–304.

56. Greiner M, Anagnostopoulos A, Pohl D, Zbinden R, Zbinden A. A rare case of severe gastroenteritis caused by *Aeromonas hydrophila* after colectomy in a patient with anti-Hu syndrome: a case report. BMC Infect Dis. 2021;21(1):1097.

57. Kumar KJ, Kumar GS. Cholera-like illness due to *Aeromonas caviae*. Indian Pediatr. 2013;50(10):969–70.

58. Marques PH, Prado LCDS, Felice AG, Rodrigues TCV, Pereira UP, Jaiswal AK, et al. Insights into the *Vibrio* genus: A One Health perspective from host adaptability and antibiotic resistance to *in silico* identification of drug targets. Antibiotics (Basel). 2022;11(10).

59. Lin H, Yu M, Wang X, Zhang XH. Comparative genomic analysis reveals the evolution and environmental adaptation strategies of vibrios. BMC Genomics. 2018;19(1):135.

60. Sinkovics JG. Horizontal gene transfers with or without cell fusions in all categories of the living matter. Adv Exp Med Biol. 2011;714:5–89.

61. Igbinosa EO, Okoh AI. *Vibrio fluvialis*: an unusual enteric pathogen of increasing public health concern. Int J Environ Res Public Health. 2010;7(10):3628–43.

62. Bhandari M, Rathnayake IU, Huygens F, Jennison AV. Clinical and Environmental *Vibrio cholerae* non-O1, non-O139 strains from Australia have similar virulence and antimicrobial resistance gene profiles. Microbiol Spectr. 2023;11(1):e0263122.

63. Shapiro BJ, Levade I, Kovacikova G, Taylor RK, Almagro-Moreno S. Origins of pandemic *Vibrio cholerae* from environmental gene pools. Nat Microbiol. 2016;2:16240.

64. Alam M, Hasan NA, Sadique A, Bhuiyan NA, Ahmed KU, Nusrin S, et al. Seasonal cholera caused by *Vibrio cholerae* serogroups O1 and O139 in the coastal aquatic environment of Bangladesh. Appl Environ Microbiol. 2006;72(6):4096–104.

65. Alam M, Sultana M, Nair GB, Sack RB, Sack DA, Siddique AK, et al. Toxigenic *Vibrio cholerae* in the aquatic environment of Mathbaria, Bangladesh. Appl Environ Microbiol. 2006;72(4):2849–55.

66. Das B, Verma J, Kumar P, Ghosh A, Ramamurthy T. Antibiotic resistance in *Vibrio cholerae*: Understanding the ecology of resistance genes and mechanisms. Vaccine. 2020;38 Suppl 1:A83–A92.

