## Supplementary material for "Genomic Diversity and Antimicrobial Resistance of *Vibrio cholerae* Isolates from Africa: A PulseNet Africa Initiative Using Nanopore Sequencing to Enhance Genomic Surveillance": File S1

**1. Methods Used for Isolation and Characterisation of *Vibrio* Isolates**

**Isolation Method:**

For stool samples: Stool samples from suspected cholera patients were collected in clean containers and processed following standard microbiological methods. Samples were plated on Thiosulfate Citrate Bile Salts Sucrose (TCBS) agar (Oxoid, UK), both directly and after 6 hours of incubation in alkaline peptone water (APW). Plates were incubated at 37°C for 18–24 hours. Single yellow colonies indicative of *V. cholerae* were selected, sub-cultured on Mueller-Hinton agar (Oxoid, UK), and incubated at 37°C for 24 hours.

For wastewater samples: 1 ml of enriched wastewater in 9 ml of Eau Peptonnée Alcaline (EPA) was incubated at 37°C for 24 hours. Samples were plated on TCBS agar, nutritive agar, and enriched in alkaline peptone and Mueller Kauffman broths. Yellow, "Chinese hat"-shaped colonies on TCBS agar were sub-cultured on Nutritive Agar (GNA) and incubated at 37°C for 24 hours, followed by Gram staining and biochemical tests (oxidase, motility, and Leminor's test).

**Characterization Techniques:**

An oxidase test was performed on colonies grown on Mueller-Hinton agar. Oxidase-positive colonies were serotyped using O1 polyvalent, O1 Inaba, O1 Ogawa, and O139 antisera (Mast Diagnostics, UK). Colonies agglutinating with O1 polyvalent and specific antisera were designated as *V. cholerae* O1 Inaba or Ogawa. Biochemical tests confirmed characteristics typical of *Vibrio* spp., such as Gram-negative curved bacilli, oxidase positivity, and other traits: citrate +, urea -, indole -, gas +, H₂S -, LDC +, and LDA -.

Serogrouping and serotyping were confirmed using Vibrio cholerae O1 antiserum (Bio-Rad, France).

**2. Epidemiological Data on Outbreaks**

**2023/2024 Cholera Outbreak in Zambia:**

**Total Cases:** 23,356 cases with 740 deaths

**Outbreak Description:** The index case for the 2023/2024 cholera outbreak was reported in Lusaka Province in October 2023, a cholera-prone region. From October 2023 to June 2024, cases were reported in 72 of 116 districts across all 10 provinces of Zambia. In January 2024, an Oral Cholera Vaccine (OCV) campaign targeted high-burden areas, achieving 99% coverage (1,870,375/1,888,112 of the target population) [2, 3].

**2014–2015 Cholera Outbreak in Côte d'Ivoire:**

The epidemic began on October 8, 2014, on an island south of Abidjan and spread inland, reaching towns over 200 km away. The last cases were reported in February 2015 in interior health districts. A total of 456 suspected cases were recorded, with 109 confirmed as *Vibrio cholerae* O1 El Tor Ogawa.

**Confirmed Cases by Month**

| Month | Year | Number of *Vibrio* isolates |
| --- | --- | --- |
| October | 2014 | 15 |
| November | 2014 | 37 |
| December | 2014 | 27 |
| January | 2015 | 28 |
| February | 2015 | 3 |

***V. cholerae* ST69 - 2023 Outbreak in South Africa**

From 1 February to 11 October 2023, a cumulative total of 1304 cholera cases were reported in seven provinces, of which 200 were laboratory-confirmed. Laboratory-confirmed cases of cholera have been identified in six provinces to date. Gauteng Province accounts for most of the cases at 88% (177/200) of the total cases followed by Free State Province with 6% (11/200), North West with 3% (6/200), Limpopo with 2% (4/200), KwaZulu-Natal with 0% (1/200) and Mpumalanga with 0% (1/200) of the total laboratory-confirmed cases. Cases have been diagnosed at both public (94%; 187/200) and private (6%; 13/200) laboratories.

The ages range from 1 to 91 years, with an average age of 39 years and a median age of 41 years. Females accounted for 51% (102/200) of the laboratory-confirmed cases. Age group 41-50 years accounted for 23% (46/200) of cases; followed by 31-40 years at 16% (33/200) and >60 years at 14% (27/200).

***V. cholerae* ST75 - 2024 Outbreak in South Africa**

From 1 December 2023 to 18 April 2024, a cumulative total of 101 suspected cholera cases were notified through the Notifiable Medical Condition (NMC) system from the Limpopo Province, of which 11 were laboratory-confirmed. Of the 11 confirmed cases 3 were imported cases with travel history to Zimbabwe (No definite history of travel or contact with a confirmed cases could be established for the other confirmed cases).

All confirmed cases have been diagnosed at public (11/11; 100%) laboratories.

The ages range from 12 to 56 years, with a median age of 31 years. Males accounted for 82% (9/11).

**References**

1. Centers for Disease Control and Prevention. Laboratory methods for the diagnosis of epidemic dysentery and cholera. In: Laboratory Methods for the Diagnosis of Epidemic Dysentery and Cholera. CDC: Atlanta, GA, 1999.
2. Zambia National Public Health Institute. <https://w2.znphi.co.zm/resources/>
3. UNICEF Zambia. Flash Update: Cholera, March 7, 2024. <https://www.unicef.org/media/153511/file/Zambia-Flash-Update-Cholera-07-March-2024.pdf>
